## Supplementary material for "ProtTrans: Towards Cracking the Language of Life’s Code Through Self-Supervised Learning": SOM

### Supplementary Online Material (SOM)

#### -1.1 Datasets

**Language model corpora.** Here, we show more details on the differences between the different corpora used for protein LM pre-training. Towards this end, we compare the number of sequences, residues as well as the amino acid distribution in UniRef50, UniRef100 and BFD. We also compare the storage size required after converting the protein sequences in each of the databases to tensorflow records (SOM Fig. 10).

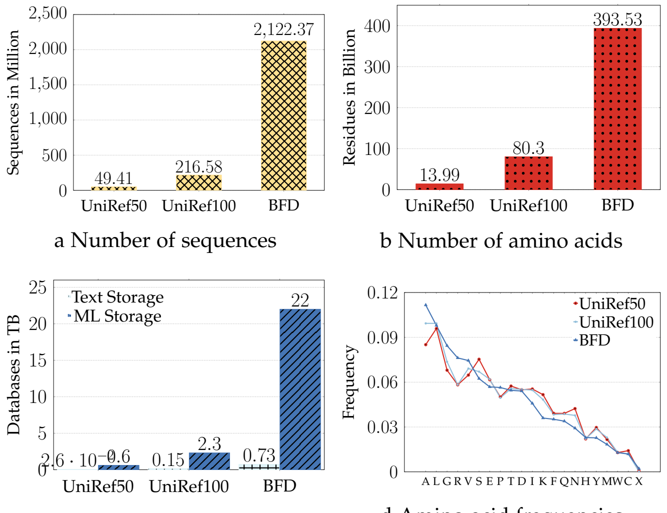


a Number of sequences

b Number of amino acids

c Storage size

d Amino acid frequencies

Fig. 10: Large Scale Dataset Training: here we compare the three datasets that were used in this study for language modelling (UniRef50, UniRef100, BFD). a) shows the number of sequences in each dataset in millions. (b) shows the number of residues/tokens in each dataset in billions. (c) shows size of each dataset raw text files as well as after converting to tensors in terabytes. (d) shows the frequency of each amino-acid/token in the each dataset

**Secondary structure data sets.** We also give a detailed overview of the three- and eight-state secondary structure class distribution for the NetSurfP-2.0 [15] training set (Train), the three test sets used in NetSurfP-2.0 (CASP12 [49], TS115 [48], CB513 [47]) as well as our new test set (NEW364) in SOM Tables 6, 5.

| Dataset | H | E | - | S | T | G | B | I |
|---|---|---|---|---|---|---|---|---|
| CASP12 | 1989 | 1416 | 1400 | 668 | 633 | 215 | 62 | 37 |
| TS115 | 10434 | 5085 | 5395 | 2210 | 2875 | 1033 | 295 | 174 |
| CB513 | 25559 | 17585 | 17713 | 8211 | 9711 | 3074 | 1105 | 469 |
| NEW364 | 26182 | 16563 | 14911 | 6233 | 7923 | 2732 | 797 | 15 |
| Train | 888175 | 559370 | 504272 | 209840 | 282561 | 99799 | 26420 | 14178 |

TABLE 5: Class distribution 8-state secondary structure - A detailed overview of the class distribution for the secondary structure datasets in 8-states is given. We compare the original NetSurfP-2.0 training set (Train), the corresponding validation datasets (CASP12, TS115, CB513) and our new test set (NEW364).

#### -1.2 Step 1: Protein LMs unsupervised

The protein LMs trained here can also be used without any supervised training either by performing a qualitative analysis of the embedding space by projecting the high-dimensional representations for a set of proteins to 2D using e.g. t-SNE. Towards this end, proteins from SCOPe (v2.07 clustered at 40% PIDE) were used as proxy for

| Dataset | Proteins | Helix | Strand | Irregular |
|---|---|---|---|---|
| CASP12 | 21 | 1478 | 2241 | 2701 |
| TS115 | 115 | 5380 | 11641 | 10480 |
| CB513 | 511 | 18690 | 29102 | 35635 |
| NEW364 | 363 | 17360 | 28929 | 29067 |
| Train | 10796 | 585790 | 1002152 | 996673 |

TABLE 6: Class distribution 3-state secondary structure - An overview of the class distribution for 3-state secondary structure data sets used in this work is given. We compare the original NetSurfP-2.0 training set (Train), the corresponding validation datasets (CASP12, TS115, CB513) and our new test set (NEW364).

protein structure (dubbed *SCOPe* in SOM Fig14-19), the three kingdoms of life (dubbed *Lineage*) and Virus as well as protein function (dubbed *Function*). For the functional classification we used only the subset of experimentally annotated EC (Enzyme Commission [5]) numbers removing all proteins that were not mappable. Protein function was also analyzed from an orthogonal angle using a different, 30%PIDE non-redundant, protein set [16] that allowed to analyze whether or not the embeddings captured aspects of the *cellular compartment; dubbed Localization in SOM Fig14-19* and membrane-association(dubbed *Membrane vs Soluble*). Additionally, we visualized the attention mechanism for a single protein to highlight the different ways and scales that Transformers offer to analyze proteins.

**Visual embedding space analysis -** Using t-SNE projections, the information content stored within the novel embeddings was qualitatively assessed on various levels, ranging from bio-physical and bio-chemical properties of single amino acids over different aspects of protein function (E.C. numbers, subcellular localization and membrane-boundness) to the level of kingdoms of life, i.e. Eukaryota, Bacteria and Archaea (for completeness here also including Viruses). We have visualized those different protein modalities for a subset of our language models, i.e. for ProtT5-U50 (SOM Fig. 14D), ProtBERT-BFD (SOM Fig. 15D), ProtBERT (SOM Fig. 16D), ProtAlbert (SOM Fig. 17D), ProtXLNet (SOM Fig. 18D) and ProtTXL (SOM Fig. 19D). When interpreting visual entropy in those figures as a proxy for the information content learnt by the models, we can observe a similar trend than for the supervised comparison, i.e. ProtT5 seems to learn on all levels a better clustering than ProtBERT.

**Attention mechanism visualization -** In line with the idea of explainable AI which tries to move away from the black-box stereotype of neural networks towards understanding why a model made a certain prediction, the analysis of the attention mechanism [70] that is at the core of each Transformer model [10] allows, to a certain extent [71], to draw first conclusions about the inner workings and the resulting predictions of Transformers. Applied to protein sequences, it was shown that the attention mechanism of Transformers can be used to predict contacts between residue pairs that are close in 3D space but far apart in sequence space [72]. Here, we visualize the attention weights [73] of one of the Transformers trained here (ProtAlbert) to analyze the structural motif of a zinc-binding domain (SOM Fig. 11). This structural motif is crucial for DNA and RNA binding across a multitude of organisms. In order to coordinate the overall fold of zinc-fingers, the binding of four specific residues, usually two Cysteines and two Histidines, to a zinc-ion is crucial. Due to their importance for the correct functioning of this protein, these residues are well conserved across most organisms with zinc-finger domains. The visual analysis of the attention triggered by the first 33 residues of an exemplary zinc-finger (PDB: 1A1L [101]) confirms that one of the attention heads in the fifth layer of ProtAlbert mostly attends to the four residues involved in coordinating the zinc-binding which indicates that the model could have learnt to pick up the signal of this, relatively frequently occurring, structural motif. Such an analysis could allow for a cheap and fast analysis of single proteins without a) needing large labeled datasets for supervised training and b) being less influenced by the experimental bias in today's labeled databases which focus mostly on model organisms with applications to biotechnology.

##### -1.3 Supervised Learning

**Supervised architecture comparison.** In this section we compared a) different choices for the supervised network that we use to evaluate the secondary structure prediction performance of the language models trained here (Table 7), b) performance of our ProtTrans models on established datasets to simplify comparability to existing prediction methods (SOM Tables 9, 8) and c) different choices for pooling embeddings of variable length protein sequence embeddings to a fixed-size representation that can be used for classifying whole protein sequences (Table 10) and

**Supervised architecture comparison.** Different architecture choices for the supervised network used to make predictions on the token-level are possible , i.e., various networks can be used to predict secondary structure for each amino acid in a protein. Towards this end, we used embeddings derived from ProtBERT-BFD to evaluate one linear classifier (logistic regression) and three non-linear classifiers (FNN, CNN and LSTM) on secondary structure prediction performance in three- and eight-states using the two hardest test sets, i.e., CASP12 and our new test set (SOM Table 7). This analysis revealed that even a logistic regression achieves competitive performance, indicating that secondary structure information is readily available from ProtBERT embeddings. Adding non-linearity without considering neighboring token embeddings (FNN) improves prediction performance and allowing the supervised network to harness local neighboring information (CNN) improves results further. However, allowing the supervised network to learn more long-range information (LSTM) only lead to an insignificant improvement for one out of four benchmarks (Q8(CASP12)) while adding more computational complexity due to the sequential nature of LSTMs. This shows that a) Transformer models already capture most of the long-range information removing the necessity to apply LSTMs, b) applying CNNs instead of FNNs improves performance slightly, potentially due to the inductive local bias of CNNs that fits well to the local nature of secondary structure. Therefore, we used a CNN architecture to evaluate the secondary structure prediction performance of all language models trained here.

**Per-residue (token level) prediction of secondary structure.** All models were evaluated using standard measures for performance (Q3/Q8: three/eight-state per-residue accuracy, i.e. percentage of residues predicted correctly in either of the 3/8 states) on standard datasets (CASP12, TS115, CB513) and a novel, highly non-redundant test set (NEW364). While TS115 and CB513 might overestimate performance because they allowed for more redundancy, we added Q3 on TS115 and CB513 to SOM Table 9 to ease comparability to existing approaches. As Q8 largely confirmed the trend observed for Q3, we added Q8 for all sets to SOM Table 8.

| Model | Q3(CASP12) | Q3(NEW364) | Q8(CASP12) | Q8(NEW364) |
|---|---|---|---|---|
| CNN | **76.1** | **81.1** | 65.2 | **70.3** |
| FNN | 75.3 | 80.0 | 63.6 | 68.9 |
| LSTM | **76.1** | 80.9 | **65.6** | 70.0 |
| LogReg | 74.3 | 79.3 | 63.4 | 68.1 |

TABLE 7: Architecture choice - We used one of our language models (ProtBERT-BFD) to compare different choices for the supervised network that we train to make predictions on the level of single amino acids, i.e, we compared a CNN, FNN, LSTM, and a logistic regression on 3- and 8-state secondary structure prediction on two different test sets (CASP12 and our new test set). Overall, the CNN provides the best performance while being computationally more efficient than the LSTM which reaches a similar performance. Despite its lower expressive power, even the logistic regression achieves competitive performance highlighting that the language models introduced here already learnt aspects of secondary structure during pre-training.

| Dataset | CASP12 | TS115 | CB513 | NEW364 |
|---|---|---|---|---|
| DeepProtVec | 49.7 | 54.4 | 48.9 | 53.3 |
| ProtTXL* | 58.5 | 63.3 | 58.9 | 61.0 |
| ProtTXL-BFD* | 58.6 | 63.3 | 58.8 | 60.9 |
| DeepSeqVec | 61.0 | 67.2 | 62.7 | 64.8 |
| ProtXLNet* | 61.6 | 68.6 | 63.1 | 65.6 |
| ProtElectra* | 60.9 | 69.1 | 64.7 | 66.9 |
| ProtAlbert* | 62.1 | 69.9 | 64.9 | 66.9 |
| ProtBert* | 63.3 | 71.5 | 66.6 | 68.9 |
| ProtBert-BFD* | 65.1 | 73.3 | 69.6 | 70.4 |
| ESM-1b | 66.0 | 73.4 | 70.2 | 71.3 |
| ProtT5-XXL-BFD* | 66.0 | 73.4 | 69.8 | 70.5 |
| ProtT5-XL-BFD* | 66.4 | 74.2 | 71.0 | 71.2 |
| ProtT5-XXL-U50* | 68.1 | 75.1 | 71.6 | 72.5 |
| NetSurfP-2.0 | 70.3 | 75.0 | 72.3 | 73.9 |
| ProtT5-XL-U50* | **70.5** | **77.1** | **74.5** | **74.5** |

TABLE 8: 8-state secondary structure prediction performance (Q8) - A detailed overview of the accuracy for correctly predicting secondary structure in 8-states is given here. We compare the performance of all language models trained here (marked with star), one word2vec-based approach (DeepProtVec), one LSTM-based (DeepSeqVec), one Transformer-based (ESM-1b) and one of the current state-of-the-art approaches that utilizes evolutionary information (NetSurfP-2.0) on three existing validation datasets (CASP12, TS115, CB513) and our new test set (NEW364). Standard errors were computed using bootstrapping: CASP12=±1.9, TS115=±1.0,CB513=±0.6,NEW364=±0.7.

Additionally, we added a detailed comparison of the 3-state secondary structure prediction performance (Q3) on NEW364 between NetSurfP-2.0 and our best performing protein LM (ProtT5-XL-U50; SOM Fig. 12). Each dot in the scatter plot reflects the Q3 achieved by NetSurfP2.0 and ProtT5-XL-U50 for the same protein. While using only single protein sequences, ProtT5-XL-U50 outperforms NetSurfP-2.0 for 57% (208 out of 364) proteins.

**MSA generation for Neff analysis.** The number of effective sequences (Neff) was computed using MSAs generated

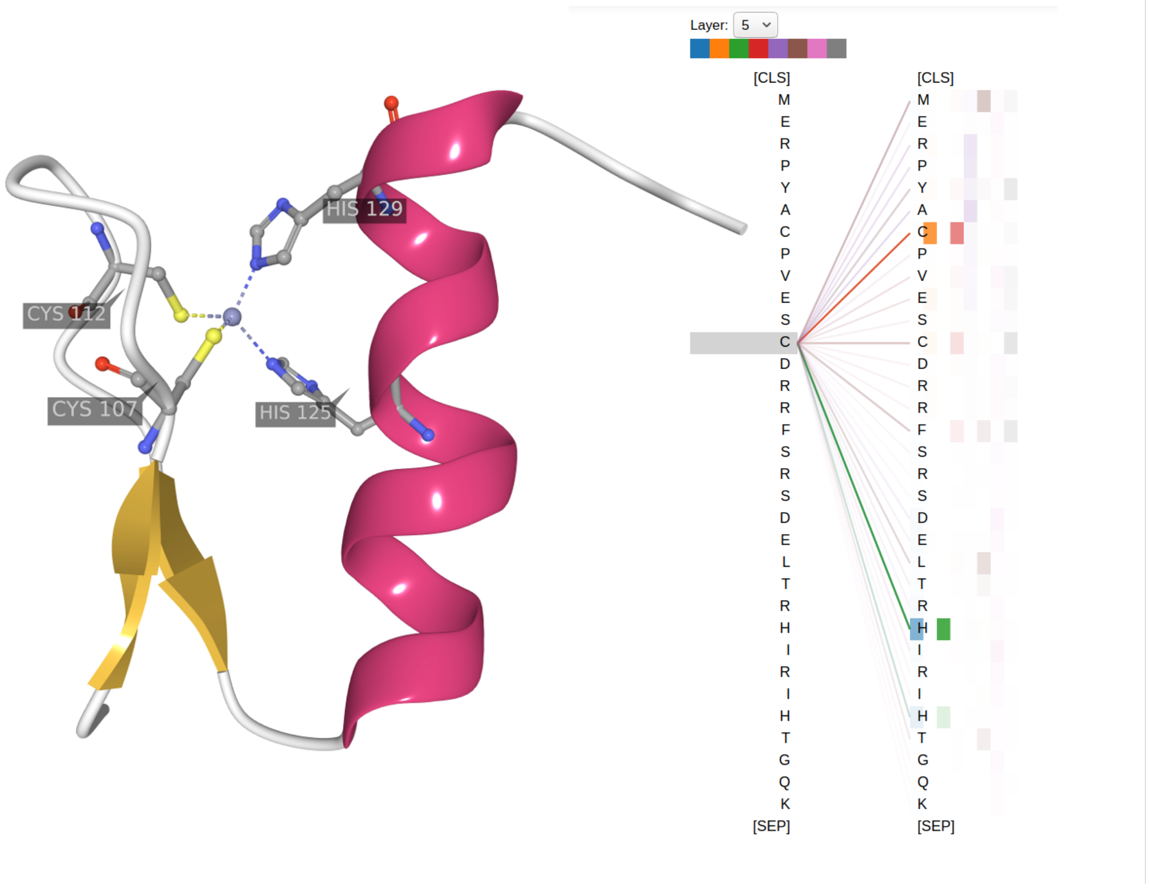


Fig. 11: Attention visualization - Here, we show how the inner workings of the Transformer's attention heads can be used to analyze single proteins in more detail. Each attention head performs internally an all-against-all comparison to compute weighted sums for each token over all other tokens in the sequence, i.e., in our case all residues in a protein are compared to each other. High scores indicate that the model learnt to put more weight on certain residue-pairs. Those scores can be indicative of a more fundamental biological truth, e.g. it was shown that some of these scores correlate with residue contacts [72]. Here, we have used bioviz [73] to visualize a protein structural motif that is crucial for DNA or RNA binding, i.e., the structure of the first 33 residues of a zinc-finger binding domain (PDB: 1A1L [101]) is shown on the left. The four residues that coordinate the zinc-binding in order to stabilize the fold are highlighted (C107, C112, H125, H129). On the right, we show a subset of the attention scores of one of our language models (ProtAlbert) for the same sequence. When visualizing the attention weights for C112, we observed that some of the attention heads in the fifth layer of ProtAlbert learnt to detect the Zinc-finger motif that is separate in sequence space but close in structure space. The attention weight is given by line thickness which indicates that the attention of residue C112 mostly attends to the three other residues (C107, H125, H129) involved in zinc-binding coordination. This shows exemplary how pre-trained Transformer models can be used for cheap and fast analysis that might open the door for novel ways of hypothesis generation.

via MMSeqs2 [51]. Toward this end, we searched all proteins in NEW364 against UniRef50 [41] using 3 iterations (–*num_iterations 3*). The UniRef50 hits were expanded by their UniRef100 cluster members (*expandaln*), effectively searching UniRef100 while reaching approximately the speed of searching UniRef50. The resulting MSA was clustered at 62% PIDE (*filterresult –max-seq-id 0.62*) to compute Neff.

**Pooling comparison.** When classifying a whole protein sequence instead of single tokens (protein-level predictions), one option is to pool token-level embeddings to derive a fixed-length protein representation from variable-length proteins. See Fig. 1 for an illustration of this process. Here, we compared four parameter-free pooling choices, i.e., min-, max-, mean-pooling (also called global average pooling) as well as a concatenation of those three. Classification accuracy when using those representations as an input for a FNN to differentiate between a) ten different subcellular localizations and b) membrane-bound and water soluble proteins is reported in Table 10. Min- and max-pooling perform significantly worse than mean-pooling. The concatenation of the three pooling strategies reaches the same performance as mean-pooling for differentiating between membrane-bound and water soluble proteins but falls significantly short on the classification of subcellular localization. One possible explanation for the latter observation is that the magnitudes of the concatenated vectors differed too much (no normaliziation was applied) while the dataset is too small for the network to learn to ignore the less informative inputs. Therefor, we stick to the mean-pooling strategy when comparing the language models trained here on classifying whole protein sequences.

| *Dataset* | *TS115* | *CB513* |
|---|---|---|
| DeepProtVec | 66.5 | 63.7 |
| ProtTXL* | 75.3 | 73.7 |
| ProtTXL-BFD* | 75.3 | 73.5 |
| DeepSeqVec | 79.0 | 77.0 |
| ProtXLNet* | 80.5 | 77.9 |
| ProtElectra* | 81.2 | 79.2 |
| ProtAlbert* | 81.9 | 79.3 |
| ProtBert* | 82.9 | 80.9 |
| ProtBert-BFD* | 83.8 | 82.5 |
| ESM-1b | 84.8 | 83.9 |
| ProtT5-XXL-BFD* | 84.6 | 83.2 |
| ProtT5-XL-BFD* | 84.8 | 83.9 |
| ProtT5-XXL-U50* | 85.6 | 84.6 |
| ProtT5-XL-U50* | **86.9** | **86.2** |
| NetSurfP-2.0 | 85.7 | 85.4 |

TABLE 9: The three-state accuracy (Q3) for the per-residue/token-level secondary structure prediction (percentage of residues/tokens correctly predicted in either of three states: helix, strand, or other) for all protein language models (LMs) trained for this work (marked by star) along with other LMs, namely one word2vec-based approach (DeepProtVec), one LSTM (DeepSeqVec), one transformer (ESM-1b) and one of the current state-of-the-art methods (NetSurfP-2.0) that uses evolutionary information (EI)/multiple sequence alignments (MSAs). Values were compiled for two datasets: one because it is a standard in the field (CASP12, results for two other standard data sets - TS115 and CB513 - in Table 6), the other because it is larger and less redundant (dubbed NEW364 introduced here). Standard errors were computed using bootstrapping: CASP12=±1.6%, NEW364=±0.5%. Highest values in each column marked in boldface.

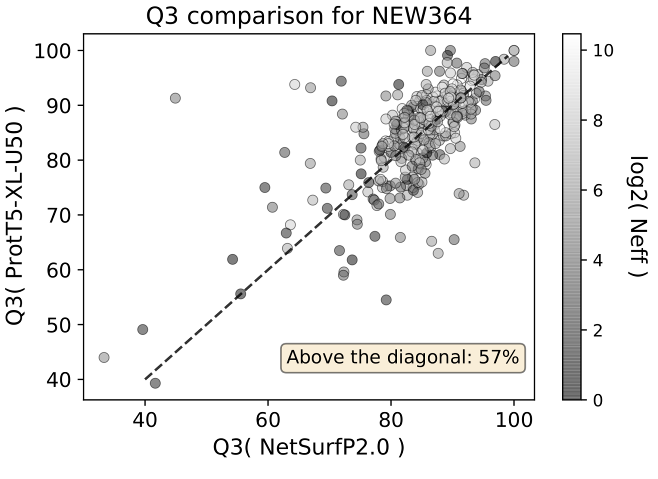


Fig. 12: Detailed Q3 comparison - We compared the 3-state secondary structure prediction performance (Q3) between one state-of-the-art method using evolutionary information (NetSurfP-2.0) and the best performing protein LM trained here (ProtT5-XL-U50) using our new test set (NEW364, see Methods). Dots resemble the Q3 achieved by each method for the same protein. Dots above the line indicate that the performance of our LM was better while dots below the diagonal show that NetSurfP-2.0 achieved higher performance. In total more than half of the proteins (57% or 208 out of 364) were predicted with higher accuracy by our method while using only single protein sequences.

| Pooling Strategy | Localization (Q10) | Membrane (Q2) |
|---|---|---|
| Min | 59 | 86 |
| Max | 60 | 85 |
| Mean | **74** | **89** |
| Concat | 64 | **89** |

TABLE 10: Pooling choice - One of our language models (ProtBERT-BFD) was used to compare different choices for pooling variable-length, token-level embeddings of a protein to a single, fixed-size protein-level representation, i.e., we compared the classification accuracy for differentiating 10 subcellular localizations and whether a protein is membrane-bound or water soluble. Global average pooling (Mean) outperforms min- or max-pooling and also the concatenation of all three pooling strategies (Concat) falls significantly short when classifying ten subcellular localizations while reaching the same performance for the classification of membrane-bound proteins.

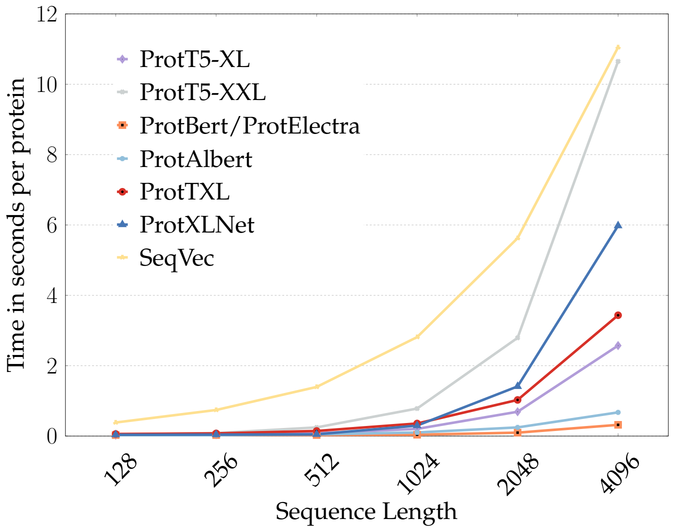


Fig. 13: Inference speed depends on sequence length: The effect of protein sequence length on the inference time of the protein LMs trained here and a previously published LM (SeqVec) were compared using a Nvidia Quadro RTX 8000 with 48GB memory using half precision (batch-size=1). Longer proteins take disproportionate long to embed for all language models. In particular, SeqVec was affected due to the sequential nature of the LSTMs used this LM, followed by ProtT5-XXL due to it is large number of parameters (5.5B for the encoder). In general, Transformer-based models require more inference time for longer for proteins because the attention maps that need to be computed square with sequence length. However, in contrast to LSTMs, the computation of attention can be parallelized, resulting in overall lower inference time for long proteins when using transformer-based LMs.

##### -1.4 Protein LM inference speed

In this section we compare the effect of sequence length on the time needed to extract features from the differen protein LMs trained here (SOM Fig. 13). Additionally, we analyse the cross-effect of sequence length and batch-size (SOM Table 11). Further, we add technical details to the human proteome benchmark shown in Fig. 9.

**Sequence length effect.** The effect of varying sequence lengths (128, 256, 512) and different batch sizes (1, 16, 32) on the inference time of the protein LMs trained here is reported in table 11. The effect of sequence length on different LM architectures (LSTM-based SeqVec and Transformer-based ProtTrans models) was visualized in figure 13. The x-axis refers to different sequence length from 128 up to 4096,

| Model | | *ProtT5-XL* | **ProtT5-XXL** | *ProtBert* | *ProtAlbert* | *ProtElectra* | *ProtXLNet* | *ProtTXL* | **SeqVec** |
|---|---|---|---|---|---|---|---|---|---|
| **Sequence Length** | **Batch Size** | | | | | | | | |
| 512 | 1 | 0.062 | 0.230 | 0.019 | 0.049 | 0.019 | 0.046 | 0.044 | 1.028 |
| | 16 | 0.054 | 0.227 | 0.013 | 0.044 | 0.13 | 0.098 | 0.054 | 0.078 |
| | 32 | 0.055 | 0.232 | 0.013 | 0.045 | 0.13 | 0.100 | 0.065 | 0.045 |
| 256 | 1 | 0.030 | 0.072 | 0.019 | 0.025 | 0.019 | 0.031 | 0.041 | 0.530 |
| | 16 | 0.015 | 0.062 | 0.005 | 0.021 | 0.005 | 0.023 | 0.012 | 0.039 |
| | 32 | 0.014 | 0.062 | 0.005 | 0.021 | 0.005 | 0.025 | 0.012 | 0.022 |
| 128 | 1 | 0.017 | 0.033 | 0.019 | 0.016 | 0.019 | 0.031 | 0.042 | 0.275 |
| | 16 | 0.006 | 0.023 | 0.003 | 0.10 | 0.003 | 0.006 | 0.004 | 0.021 |
| | 32 | 0.006 | 0.023 | 0.002 | 0.10 | 0.002 | 0.006 | 0.004 | 0.012 |
| Average | 1 | 0.036 | 0.112 | 0.019 | 0.03 | 0.019 | **0.036** | 0.042 | 0.611 |
| | 16 | 0.025 | **0.104** | 0.007 | 0.025 | 0.007 | 0.042 | **0.023** | 0.046 |
| | 32 | **0.025** | 0.106 | **0.007** | **0.025** | **0.007** | 0.044 | 0.027 | **0.026** |

TABLE 11: Comparison of inference speed: The analysis distinguished proteins of different length, as well as different batch sizes (numbers of proteins processed: 1, 16 and 32. For simplicity, no proteins longer than 512 is shown . Each test was repeated 100 times and the average time per protein was reported. The experiment was conducted using a single redNvidia Quadro RTX8000 with 48GB memory.

while the y-axis represents the time of inference in ms for a single protein with a batch size of 1 on a Nvidia Quadro RTX8000 with 48GB.

##### -1.5 Fast, proteome-wide feature-extraction.

We compared the time required to generate features between established methods (EI/MSA-based) and the proposed embeddings derived from the protein LMs trained here. Towards this end, we created embeddings for each protein in the human proteome (20,353) proteins with a median sequence length of 415 residues) using a) our protein LMs to generate embeddings and b) the fastest method available, namely MMseqs2 [51], to generate MSAs. In light of wanting to compare to the state-of-the-art secondary structure prediction method, NetSurfP-2.0 [15], we used the same parameters for the MMseqs2 search used by that method (*–num_iterations 2 –diff 2000*) and compared two databases (UniRef90 with 113M and UniRef100 with 216M proteins). All comparisons used an IntelR© XeonR© Scalable Processor "Skylake" Gold 6248 with 40 threads, SSD and 377GB main memory, while protein LMs were computed on a single NVIDIA Quadro RTX 8000 with 48GB memory using half precision and dynamic batch size based on variable protein sequence lengths. MMseqs2 was about 16 to 28-times slower than the fastest LMs (ProtElectra and ProtBert), and about 4 to 6-times slower than our best model (ProtT5) (Fig. 9). The best performing model, ProtT5-XL-U50, required on average 0.12 seconds to create embeddings for a human protein, completing the entire human proteome (all proteins in an organism) in 40 minutes. We noticed that SeqVec was the slowest model for long proteins (11 seconds for protein with 4096 residues) while ProtBert was the fastest (0.32s) for those.

##### -1.6 Additional Resources

For long-term storage the repository is also backed up through zenodo[9]. Additionally, all models are in the transformer library of *huggingface* [15][10]. Furthermore, secondary structure predictions for ProtBERT-BFD are available through the PredictProtein [25] webserver[11]. Furthermore, the bio_embeddings [17] package[12] package could be used for simplified accessibility and analysis of protein LMs and predictions of the supervised models.

9. https://zenodo.org/record/4633482
10. https://huggingface.co/Rostlab
11. https://predictprotein.org/
12. https://github.com/sacdallago/bio_embeddings/

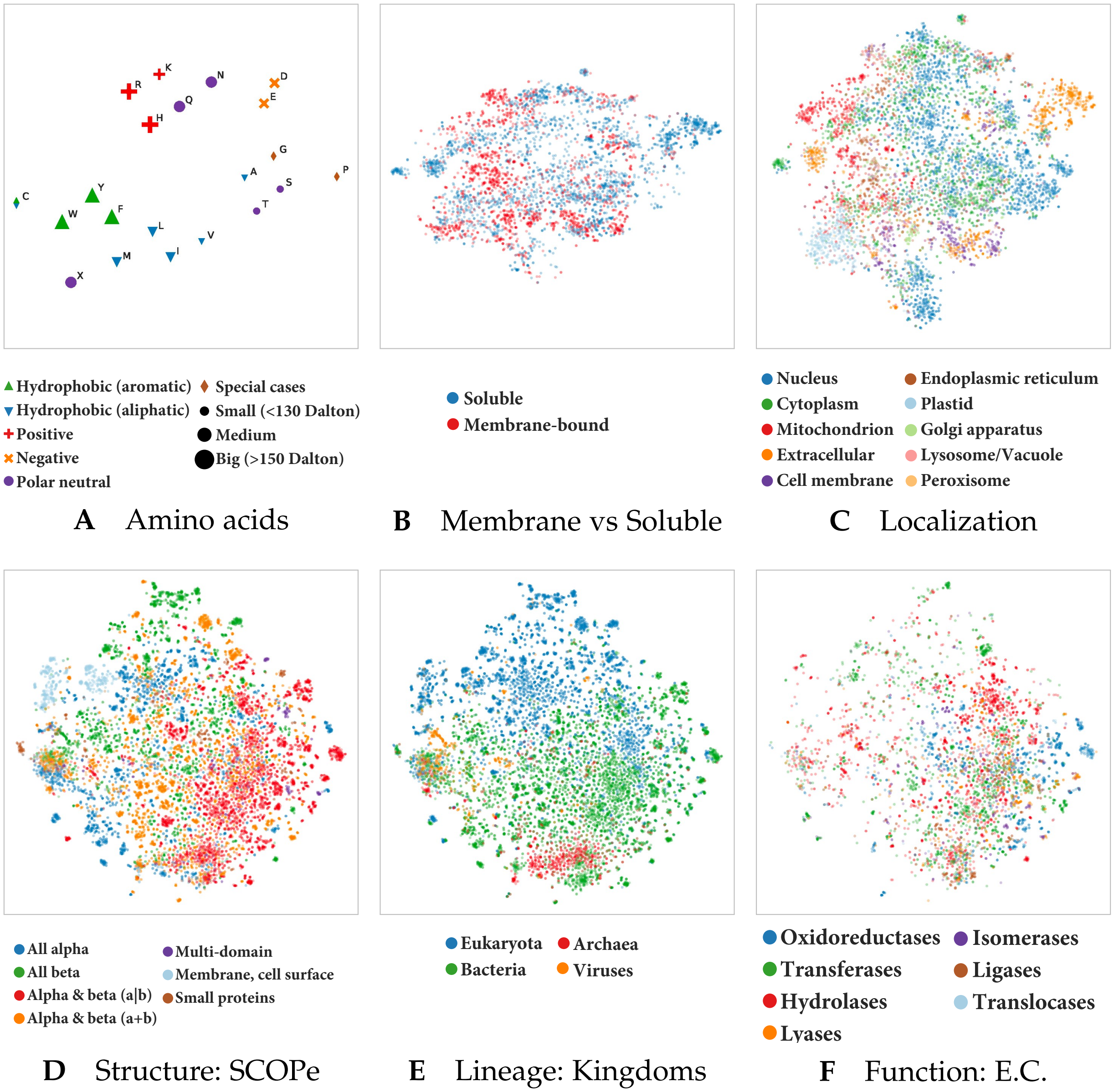


Fig. 14: Unsupervised training captures various features of proteins: We used t-SNE projections to assess which features the LMs trained here learnt to extract from proteins. Exemplarily for ProtT5-XL-U50, the best-performing model on supervised tasks, we showed that the protein LMs trained here captured biophysical- and biochemical properties of single amino acids during pre-training (Panel A). A redundancy reduced version (30%) of the DeepLoc [16] dataset was used to assess whether the LM learnt to classify proteins into membrane-bound and water-soluble (Panel B) or according to their cellular compartment (Panel C). Not all proteins in the set had annotations for both features, making Panels B and C not directly comparable. Further, a redundancy reduced version (40%) of the Structural Classification of Proteins – extended (SCOPe) database was used to assess whether ProtT5-XL captured structural (Panel D), functional (Panel F) or lineage-specific (Panel E) features of proteins without any labels. Towards this end, contextualized, fixed-size representations were generated for all proteins in both datasets by mean-pooling over the representations extracted from the last layer of ProtT5-XL (average over the length of the protein). The high-dimensional embeddings were projected to 2D using t-SNE. ProtT5-XL captured protein information on different levels: ranging from structural features as annotated in the main classes in SCOPe, over functional aspects as defined by in the Enzyme Commission (E.C.) numbers or the cellular compartment to the branch of the protein within the tree of life, without ever having been explicitly trained on any of these features. Comparing different features for the same datasets revealed that potentially heterogeneous clusters are only formed due to the multi-modal nature of proteins, e.g. the eukaryotic proteins are well separated from bacterial proteins (Panel E) but form internally multiple sub-clusters in structure space (Panel D).

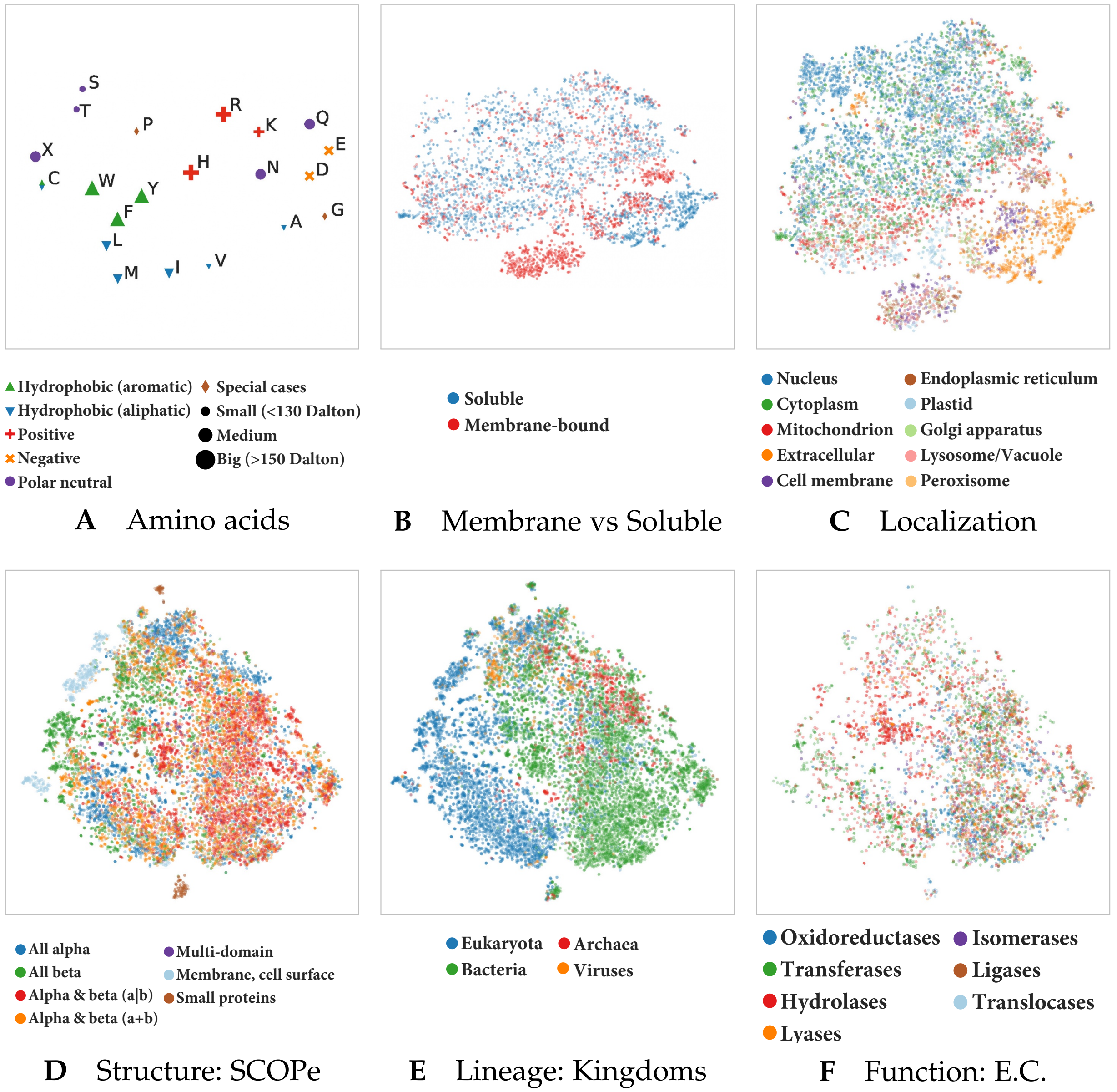


Fig. 15: Unsupervised training captures various features of proteins: We used t-SNE projections to assess which features the LMs trained here learnt to extract from proteins. Exemplarily for ProtBert-BFD, we showed that the protein LMs trained here captured biophysical- and biochemical properties of single amino acids during pre-training (Panel A). A redundancy reduced version (30%) of the DeepLoc [16] dataset was used to assess whether the LM learnt to classify proteins into membrane-bound and water-soluble (Panel B) or according to their cellular compartment (Panel C). Not all proteins in the set had annotations for both features, making Panels B and C not directly comparable. Further, a redundancy reduced version (40%) of the Structural Classification of Proteins – extended (SCOPe) database was used to assess whether ProtBert-BFD captured structural (Panel D), functional (Panel F) or lineage-specific (Panel E) features of proteins without any labels. Towards this end, contextualized, fixed-size representations were generated for all proteins in both datasets by mean-pooling over the representations extracted from the last layer of ProtBert-BFD (average over the length of the protein). The high-dimensional embeddings were projected to 2D using t-SNE. ProtBert-BFD captured protein information on different levels: ranging from structural features as annotated in the main classes in SCOPe, over functional aspects as defined by in the Enzyme Commission (E.C.) numbers or the cellular compartment to the branch of the protein within the tree of life, without ever having been explicitly trained on any of these features. Comparing different features for the same datasets revealed that potentially heterogeneous clusters are only formed due to the multi-modal nature of proteins, e.g. the eukaryotic proteins are well separated from bacterial proteins (Panel E) but form internally multiple sub-clusters in structure space (Panel D).

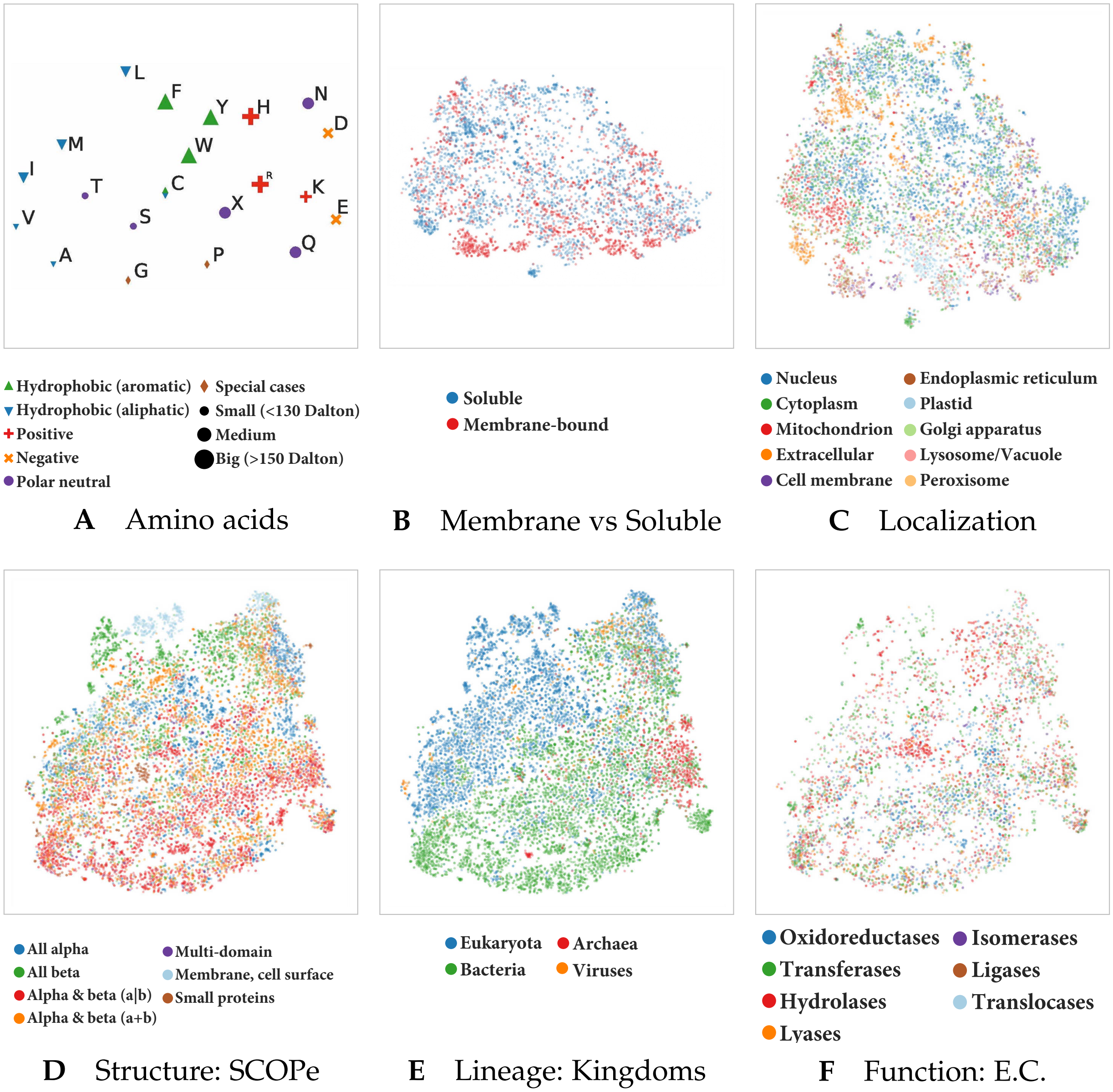


Fig. 16: Unsupervised training captures various features of proteins: We used t-SNE projections to assess which features the LMs trained here learnt to extract from proteins. Exemplarily for ProtBert, we showed that the protein language models trained here captured biophysical- and biochemical properties of single amino acids (Panel A). A redundancy reduced version (30%) of the DeepLoc ( [16]) dataset was used to assess whether ProtBert learnt to classify proteins into membrane-bound or water-soluble (Panel B) or according to the cellular compartment they appear in (Panel C). Not all proteins in the set had annotations for both features, making Panels B and C not directly comparable. Further, a redundancy reduced version (40%) of the Structural Classification of Proteins – extended (SCOPe) database was used to assess whether ProtBert captured structural (Panel D), functional (Panel F) or lineage-specific (Panel E) features of proteins without any labels. Towards this end, contextualized, fixed-size representations were generated for all proteins in both datasets by mean-pooling over the representations extracted from the last layer of ProtBert (average over the length of the protein). The high-dimensional embeddings were projected to 2D using t-SNE. ProtBert formed less dense clusters compared to the same model trained on a larger dataset (ProtBert-BFD Fig. 15).

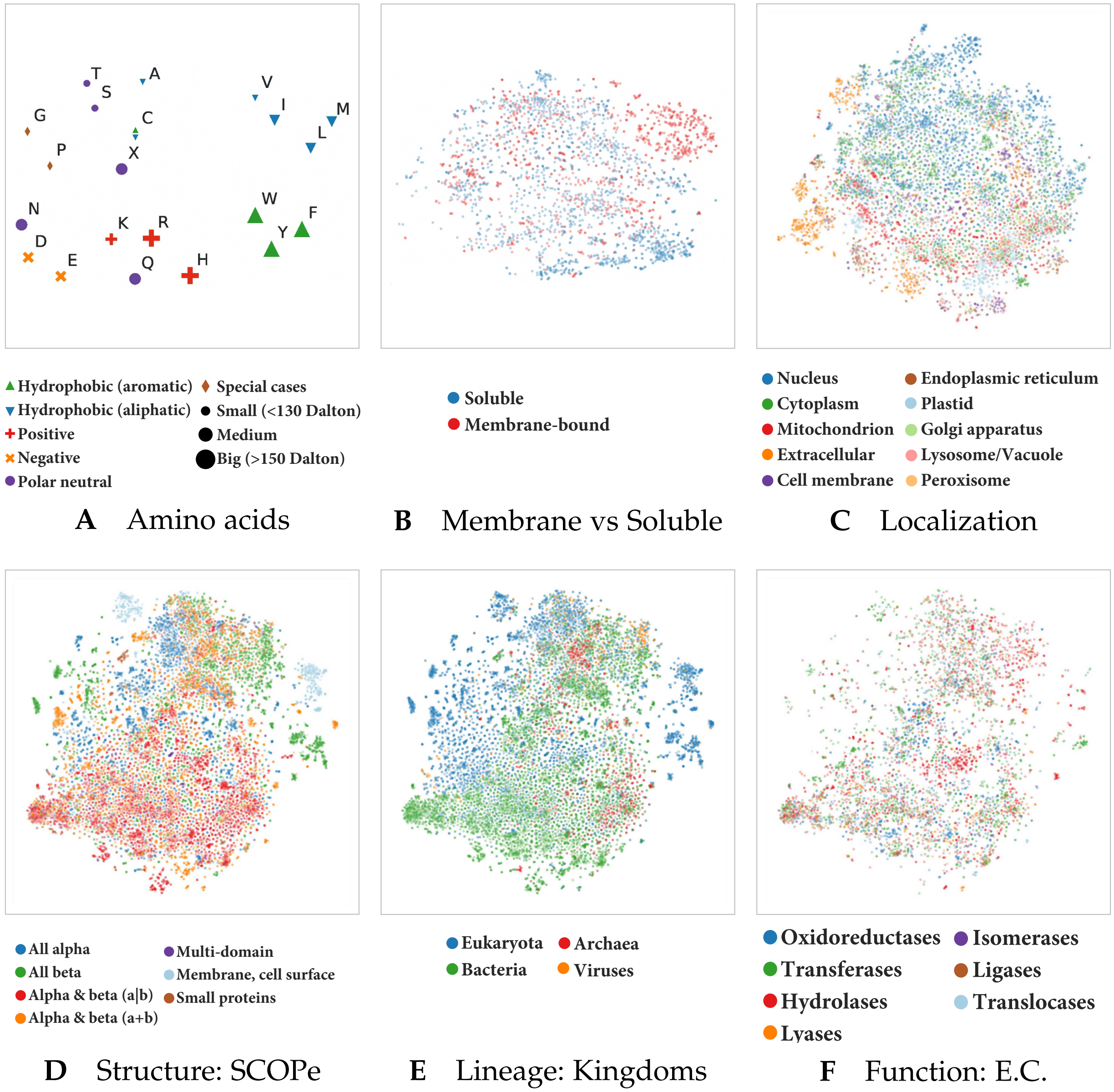


Fig. 17: Unsupervised training captures various features of proteins: We used t-SNE projections to assess which features the LMs trained here learnt to extract from proteins. Exemplarily for ProtAlbert, we showed that the protein language models trained here captured biophysical- and biochemical properties of single amino acids (Panel A). A redundancy reduced version (30%) of the DeepLoc ( [16]) dataset was used to assess whether ProtAlbert learnt to classify proteins into membrane-bound or water-soluble (Panel B) or according to the cellular compartment they appear in (Panel C). Not all proteins in the set had annotations for both features, making Panels B and C not directly comparable. Further, a redundancy reduced version (40%) of the Structural Classification of Proteins – extended (SCOPe) database was used to assess whether ProtAlbert captured structural (Panel D), functional (Panel F) or lineage-specific (Panel E) features of proteins without any labels. Towards this end, contextualized, fixed-size representations were generated for all proteins in both datasets by mean-pooling over the representations extracted from the last layer of ProtAlbert (average over the length of the protein). The high-dimensional embeddings were projected to 2D using t-SNE. Compared to the some of the other protein LMs trained here (ProtTXL 19 and ProtBert 16, ProtAlbert formed more dense clusters, especially for the projections based on the SCOPe dataset (Panels D, E and F).

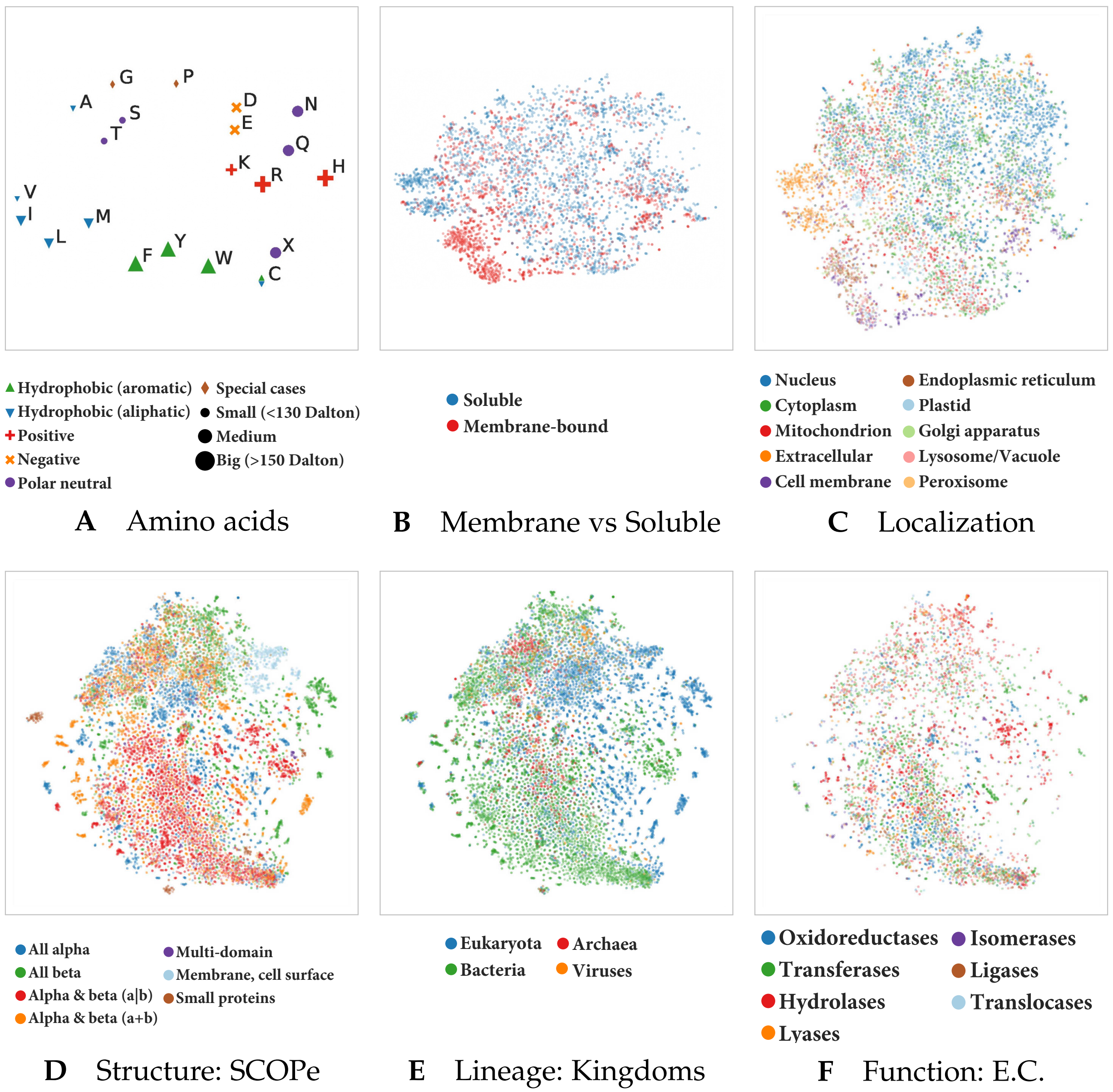


Fig. 18: Unsupervised training captures various features of proteins: We used t-SNE projections to assess which features the LMs trained here learnt to extract from proteins. Exemplarily for ProtXLNet, we showed that the protein language models trained here captured biophysical- and biochemical properties of single amino acids (Panel A). A redundancy reduced version (30%) of the DeepLoc ( [16]) dataset was used to assess whether ProtXLNet learnt to classify proteins into membrane-bound or water-soluble (Panel B) or according to the cellular compartment they appear in (Panel C). Not all proteins in the set had annotations for both features, making Panels B and C not directly comparable. Further, a redundancy reduced version (40%) of the Structural Classification of Proteins – extended (SCOPe) database was used to assess whether ProtXLNet captured structural (Panel D), functional (Panel F) or lineage-specific (Panel E) features of proteins without any labels. Towards this end, contextualized, fixed-size representations were generated for all proteins in both datasets by mean-pooling over the representations extracted from the last layer of ProtXLNet (average over the length of the protein). Compared to other protein LMs trained here, ProtXLNet learnt small coherent clusters that are scattered among the t-SNE projection. Only comparing different features for the same datasets reveals that potentially heterogenous clusters are only formed due to the mulit-modal nature of proteins, e.g. the eukaryotic proteins are well separated from bacterial proteins (Panel E but form multiple sub-clusters in structure space (Panel D). Compared to other protein LMs trained here (e.g. ProtTXL 19, ProtXLNet learnt small coherent clusters that are scattered among the t-SNE projection. Simlar to ProtBert-BFD, some of the small scattered clusters form homogeneous clusters when focusing on other aspects of proteins, e.g. some of the proteins in the heterogeneous cluster in the lower left part showing subcellular localization (Panel C) can be explained by proteins bound to the membrane (red in Panel B).

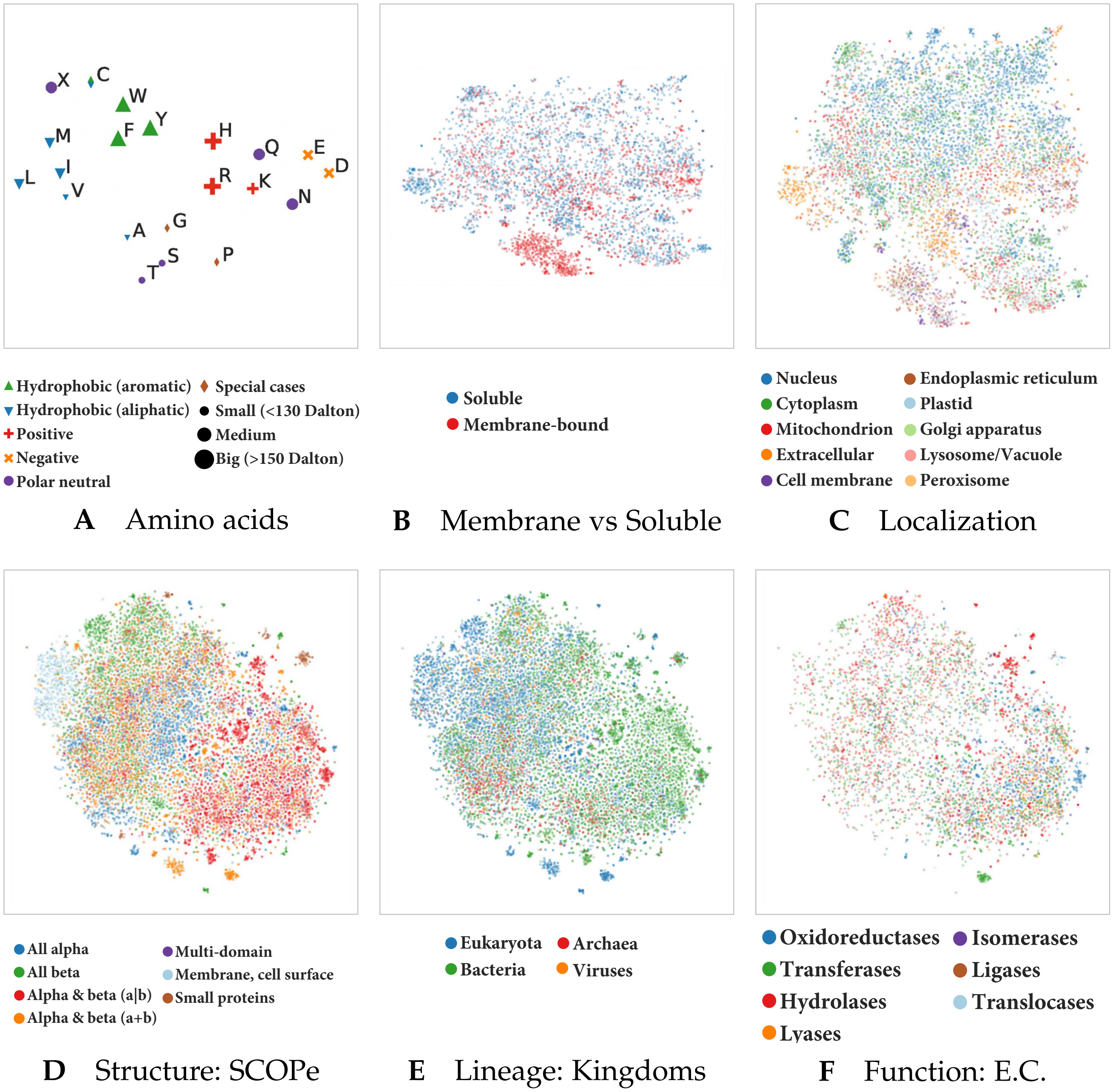


Fig. 19: Unsupervised training captures various features of proteins: We used t-SNE projections to assess which features the LMs trained here learnt to extract from proteins. Exemplarily for ProtTXL, we showed that the protein language models trained here captured biophysical- and biochemical properties of single amino acids (Panel A). A redundancy reduced version (30%) of the DeepLoc ( [16]) dataset was used to assess whether ProtTXL learnt to classify proteins into membrane-bound or water-soluble (Panel B) or according to the cellular compartment they appear in (Panel C). Not all proteins in the set had annotations for both features, making Panels B and C not directly comparable. Further, a redundancy reduced version (40%) of the Structural Classification of Proteins – extended (SCOPe) database was used to assess whether ProtTXL captured structural (Panel D), functional (Panel F) or lineage-specific (Panel E) features of proteins without any labels. Towards this end, contextualized, fixed-size representations were generated for all proteins in both datasets by mean-pooling over the representations extracted from the last layer of ProtTXL (average over the length of the protein). The high-dimensional embeddings were projected to 2D using t-SNE. While generally forming the least dense clusters compared to other LMs trained here, ProtTXL captured certain aspects about protein function (e.g. Transferases, dark green Panel F) that other LMs trained here did not capture.

### References - SOM -


[1] M. S. Klausen, M. C. Jespersen, H. Nielsen, K. K. Jensen, V. I. Jurtz, C. K. Sønderby, M. O. A. Sommer, O. Winther, M. Nielsen, B. Petersen, and P. Marcatili, "NetSurfP-2.0: Improved prediction of protein structural features by integrated deep learning," *Proteins: Structure, Function, and Bioinformatics*, vol. 87, no. 6, pp. 520–527, 2019, _eprint: https://onlinelibrary.wiley.com/doi/pdf/10.1002/prot.25674.
diction methods," *Proteins: Structure, Function, and Bioinformatics*, vol. 86, pp. 97–112, 2018.

[3] J. A. Cuff and G. J. Barton, "Evaluation and improvement of multiple sequence methods for protein secondary structure prediction," *Proteins: Structure, Function, and Bioinformatics*, vol. 34, no. 4, pp. 508–519, 1999.

[4] Y. Yang, J. Gao, J. Wang, R. Heffernan, J. Hanson, K. Paliwal, and Y. Zhou, "Sixty-five years of the long march in protein secondary structure prediction: The final stretch?" *Briefings in bioinformatics*, vol. 19, no. 3, pp. 482–494, 2018.

[5] E. C. Webb, *Enzyme Nomenclature 1992. Recommendations of the Nomenclature committee of the International Union of Biochemistry and Molecular Biology.*, 1992nd ed. New York: Academic Press, 1992.

[6] J. J. Almagro Armenteros, C. K. Sønderby, S. K. Sønderby, H. Nielsen, and O. Winther, "DeepLoc: Prediction of protein subcellular localization using deep learning," *Bioinformatics*, vol. 33, no. 21, pp. 3387–3395, Nov. 2017.

[7] D. Bahdanau, K. Cho, and Y. Bengio, "Neural Machine Translation by Jointly Learning to Align and Translate," *arXiv:1409.0473 [cs, stat]*, May 2016.

[8] A. Vaswani, N. Shazeer, N. Parmar, J. Uszkoreit, L. Jones, A. N. Gomez, t. Kaiser, and I. Polosukhin, "Attention is all you need,"

[2] L. A. Abriata, G. E. Tamò, B. Monastyrskyy, A. Kryshtafovych, and M. Dal Peraro, "Assessment of hard target modeling in CASP12 reveals an emerging role of alignment-based contact pre-

in *Advances in Neural Information Processing Systems*, 2017, pp. 5998–6008.

[9] S. Vashishth, S. Upadhyay, G. S. Tomar, and M. Faruqui, "Attention interpretability across nlp tasks," *arXiv preprint arXiv:1909.11218*, 2019.

[10] R. M. Rao, J. Meier, T. Sercu, S. Ovchinnikov, and A. Rives, "Transformer protein language models are unsupervised structure learners," *bioRxiv*, 2020. [Online]. Available: https://www.biorxiv.org/content/10.1101/2020.12.15.422761v1

[11] J. Vig, "A multiscale visualization of attention in the transformer model," 2019.

[17] C. Dallago, K. Schütze, M. Heinzinger, T. Olenyi, M. Littmann, A. X. Lu, K. K. Yang, S. Min, S. Yoon, J. T. Morton, and B. Rost, "Learned embeddings from deep learning to visualize and predict protein sets," *Current Protocols in Bioinformatics*.
